# Quantifying uncertainty in drift diffusion models of decision making under temporal dependence and parameter variability

**DOI:** 10.64898/2026.05.17.722295

**Authors:** Gabriel Riegner, Armin Schwartzman, Pamela Reinagel

## Abstract

Decision-making behavior changes over time, exhibiting temporal correlation and nonstationarity. Existing drift diffusion model (DDM) fitting methods either do not provide uncertainty quantification for parameter estimates, or rely on restrictive assumptions that decisions are independent and that parameters remain constant over time, potentially underestimating uncertainty. To address these limitations, we propose a computationally efficient method for estimating analytic uncertainties in DDM parameters that are robust to temporal dependence and unmodeled parameter variability, while explicitly modeling nonstationary variability through covariates. We apply this method to rat decision-making in a two-alternative forced-choice (2AFC) visual task, revealing dynamic decision-making states across multiple timescales. A Python implementation of the method is provided.

## 1 Introduction

### 1.1 Perceptual Decision Making in Naturalistic Settings

Perceptual decision making is the process of making decisions based on sensory information. While often studied in highly controlled experiments, in naturalistic settings these decisions are rarely stable; rather, they emerge from the interplay between multiple interleaved decision states that are richly structured in time (Roy et al., 2018, 2021; Ashwood et al., 2022; Duffy et al., 2025; Urai, 2025; Danskin et al., 2023; Perquin et al., 2023; Cochrane et al., 2023). The dominant theoretical models of these two-alternative forced-choice (2AFC) decisions are sequential sampling models, which postulate that decisions are made upon the sequential analysis of evidence (Shadlen et al., 2006; Asadpour et al., 2024). These models are supported by substantial neural evidence and underlying computational principles that describe the noisy accumulation of evidence leading to a choice (Shadlen et al., 2006; Steinemann et al., 2024; Bogacz et al., 2006; Asadpour et al., 2024; Deakin et al., 2024).

### 1.2 The Drift Diffusion Model (DDM)

In this paper, we consider the DDM, a sequential sampling model which has a long history of describing decision making by a latent diffusion process that accumulates noisy sensory evidence over time until it reaches a decision bound, simultaneously explaining which choice is made and how long it takes to make it (Stone, 1960; Ratcliff, 1978; Shadlen & Newsome, 2001). The model translates the latent diffusion process into observed behavior via parameters that together define the *decision-making state*; elsewhere in the literature this is referred to as the decision-making strategy (Ashwood et al., 2022; Roy et al., 2018, 2021). The DDM and its parameters are defined in Section 2.1.1.

### 1.3 Decision Making Over Time

The decision state is not constant; it changes over time with internal and external environments across timescales ranging from milliseconds to hours to days and beyond (Duffy et al., 2025; Urai et al., 2019; Cochrane et al., 2023; Shevinsky & Reinagel, 2019; Schumacher et al., 2023, 2025). Subjects adapt to these changing environments by adjusting their internal decision states, and these latent states in turn shape observed decision outcomes. Although standard measures of decision making often mask the temporal structure of these changes, recent approaches have begun to recover these time-varying dynamics by inferring parameter trajectories directly from behavioral data (Ashwood et al., 2022; Gunawan et al., 2022; Kucharský et al., 2021; Maggi et al., 2024). For example, subjects may alternate between discrete engaged stimulus-dependent states and disengaged stimulus-independent states (Ashwood et al., 2022; Mohammadi et al., 2025; Duffy et al., 2025) or drift slowly between continuous states (Vloeberghs et al., 2025; Roy et al., 2021, 2018), or a combination of both (Schumacher et al., 2023, 2025). As noted by Urai (2025) in the context of decision making, “there is probably no such thing as stable behavior.”

Moreover, decisions are history dependent, influenced by sequential effects including the outcomes of previous choices (Frund et al., 2014; Gao et al., 2009). Subjects use past experience to guide future choices, often integrating history information across multiple timescales (Danskin et al., 2023; Nguyen et al., 2019; Busse et al., 2011). This kind of history dependence can include effects of reinforcement learning, in which rewarded actions shape subsequent choices (Pedersen et al., 2017; Fengler et al., 2022), as well as reward-independent sequential biases (Frund et al., 2014; Gao et al., 2009). These forms of temporal structure are prevalent across perceptual decision making in humans, monkeys, and rodents (Urai et al., 2019; Busse et al., 2011; Treviño et al., 2022; Braun et al., 2018; Gupta et al., 2024). Humans may also exhibit forms of variability similar to those observed in rodents, differing only in the relative variability of different decision-making parameters (Nguyen & Reinagel, 2022; Shevinsky & Reinagel, 2019). Together, these demonstrate dynamic decision making across species, in which the past influences the present, producing trial-to-trial variability that is richly structured in time.

### 1.4 Current Estimation Methods and Their Limitations

Standard DDM fitting methods often implicitly assume that, in the absence of experimental manipulations, the decision-making state remains constant over time with fixed parameters, treating decision outcomes as independent and identically distributed (*iid*), a condition often violated in real data (Moens & Zenon, 2018; Urai, 2025; Perquin et al., 2024). When parameters are allowed to vary — e.g., the seven-parameter DDM of Ratcliff & Rouder (1998) — they do so as unstructured random variability around a stable mean, with trial order still ignored, and the possibility that systematic trends such as learning curves or circadian rhythms are linked to observable covariates is not considered (Ratcliff, 2013). Moreover, across-trial variability parameters are inherently difficult to estimate, showing poor recovery and weak test-retest reliability due to unidentifiability (Boehm et al., 2018; Lerche & Voss, 2016; Ratcliff & Childers, 2015; Ratcliff et al., 2018; Duffy et al., 2025; Lerche et al., 2017; Van Ravenzwaaij & Oberauer, 2009).

Current inference methods compound these issues in four ways.

▪ Point estimates do not always include corresponding uncertainties (Shinn et al., 2020; Boehm et al., 2018).
▪ When uncertainties are reported, they rely on asymptotic approximations that assume *iid* conditions often not met in practice (Moens & Zenon, 2018; Wiecki et al., 2013).
▪ Quantifying uncertainty typically requires computationally expensive methods such as Bayesian MCMC or resampling (Boehm et al., 2018; Moens & Zenon, 2018; Wiecki et al., 2013; Fengler et al., 2026).
▪ The robustness of inference to unmodeled parameter variability and temporal correlations has not been addressed. Under misspecification, standard errors underestimate true uncertainty, resulting in overconfident and invalid inferences (Boehm et al., 2018; Urai, 2025; Hato et al., 2025; Schumacher et al., 2025).

Together, these limitations constrain hypothesis testing and the detection of meaningful temporal changes in decision states, particularly in naturalistic settings where brain and behavior are in constant flux (Urai, 2025; Ratcliff, 2013). As a result, even when rich temporal structure is present in long behavioral time series, experimenters lack a computationally efficient way to obtain valid uncertainties and hypothesis tests for time-varying DDM parameters.

### 1.5 Proposed Solutions

We consider the case of decision making over time, which introduces dependence and complicates inference. For example, parameters may trend within a session as satiety or fatigue accumulates, or drift between sessions as subjects learn. To address this, we explicitly allow for time dependence and model misspecification in order to conduct valid inference. We additionally propose modeling DDM parameters as functions of observed covariates (such as stimulus strength, trial number, or time of day) via link functions. This captures how external factors influence each decision and provides a principled mechanism for nonstationary behavior. Systematic parameter trends that would otherwise violate stationarity are explained by the covariate model, leaving residual variability that is plausibly stable over time. Stationarity is therefore a requirement on the joint process of covariates and outcomes, not on the observed decision outcomes alone. This distinction substantially broadens the scope of experiments to which the method applies.

We introduce a computationally efficient pseudo-maximum likelihood estimation (pseudo-MLE) method for estimating DDM parameters and covariate effects, together with analytical uncertainty estimates that remain valid under model misspecification and temporal dependence. The paper is organized as follows. Section 2 formalizes the model, estimator, and large-sample theory. Numbered equations indicate key expressions referenced throughout, while the others provide intermediate steps that can be skipped without loss of continuity. We implemented all described methods in Python except where other packages are explicitly cited. Section 3 validates the method through simulations, and Section 4 demonstrates applications to rat decision making. Section 5 compares the proposed estimator with Bayesian MCMC, and Section 6 discusses implications, limitations, and future directions.

## 2 Methods

### 2.1 Notation

#### 2.1.1 Drift Diffusion Model

The DDM describes perceptual decision making in 2AFC tasks by parameterizing a latent decision process with choice and reaction time as outputs. This process accumulates noisy sensory evidence over time until it reaches one of two decision bounds, initiating the corresponding choice. A *trial* is a single instance of this process, starting with a stimulus that initiates evidence accumulation and ending with a choice and reaction time. In the present paper, the notation is kept general so that the upper and lower bounds can be mapped to any pair of alternatives (e.g., rightward vs. leftward, or correct vs. error). The standard DDM has four decision-making parameters as shown in Figure 1B.

**Figure 1:**
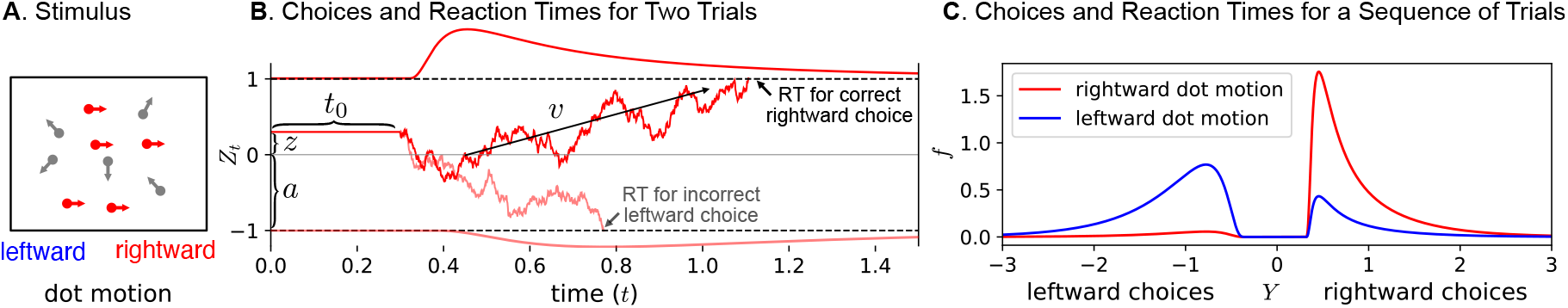
Drift Diffusion Model. The drift diffusion model describes perceptual decision making as the noisy accumulation of evidence toward one of two decision bounds. In this example, the stimulus is random dot motion and its strength is modulated by *coherence*, the percentage of dots moving in the direction of the correct choice. (A) Shows a random dot stimulus at 50% rightward coherence, indicating the correct choice is to the right. (B) Shows two realizations of the drift diffusion process *Z*_*t*_ with fixed parameters ***θ*** = (*a, z, t*_0_, *v*). The two realizations evolve as stochastic processes, crossing the upper (solid red; correct rightward) and lower (translucent red; incorrect leftward) bounds at different reaction times *RT* . The correct and error *RT* probability densities are shown above and below the panel; the correct rightward choice is more probable because *v >* 0 (proportional to rightward coherence) and *z >* 0 (rightward bias). (C) Probability densities of the composite variable *Y* encoding both the choice (negative for leftward, positive for rightward) and the reaction time (*RT* = |*Y* |) for a sequence of rightward (red) and leftward (blue) coherence trials. The proportion of correct responses for rightward trials (red) is higher than for leftward trials (blue) because *z >* 0 (rightward bias).

▪ *a >* 0 is the *decision bound* : evidence is accumulated until the decision variable crosses the upper bound at +*a* or the lower bound at −*a*, with a separation between bounds of 2*a*.
▪ *z* ∈ (−1, +1) is the *relative starting point*: the initial bias of the decision variable where sign(*z*) indicates bias toward the upper vs. lower bound and |*z*| gives the bias magnitude as a fraction of *a*.
▪ *t*_0_ ≥ 0 is the *non-decision time*: additive time in seconds capturing processing outside evidence accumulation, including sensory encoding and motor response.
▪ *v* ∈ ℝ is the *drift rate*: the expected rate of change in the decision variable per second, where sign(*v*) implies drift toward the upper vs. lower bound and |*v*| gives the evidence strength/quality.

Taken together, the vector containing the four parameters ***θ*** = (*a, z, t*_0_, *v*) defines the behavior of the latent decision process for a single trial, which we refer to as the decision-making state throughout. The decision variable *Z*_*t*_ captures evidence accumulation between the upper +*a* and lower −*a* decision bounds at time *t*, combining deterministic drift and stochastic diffusion:

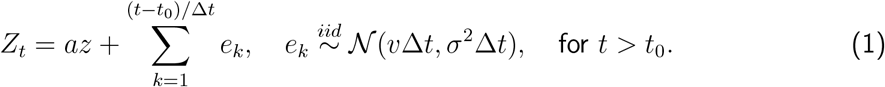

Here, *t* is the elapsed time and *t*_0_ is the non-decision time, both defined as integer multiples of the step duration Δ*t*. Evidence accumulation begins at 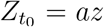 after the non-decision time *t*_0_ and proceeds as a random walk with deterministic drift. For simplicity, *t*_0_ is treated as preceding the decision process, when in reality it combines non-decision processes before and after. Each subsequent update adds an *iid* Gaussian step *e*_*k*_ with mean *v*Δ*t* and variance *σ*^2^Δ*t*; each step is random but the process drifts on average toward the upper (if *v >* 0) or lower (if *v <* 0) bound. The duration Δ*t* rescales continuous drift and diffusion rates *v* and *σ*^2^ (fixed to *σ*^2^ = 1) into discrete steps, where the summation runs over a total of (*t* −*t*_0_)*/*Δ*t* steps. When Δ*t* →0, this discrete random walk approaches a continuous drift (or Wiener) diffusion process.

#### 2.1.2. Choice and Reaction Time for a Single Trial

A decision is made at the first time *t > t*_0_ for which the decision variable *Z*_*t*_ (1) reaches either decision bound:

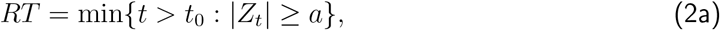

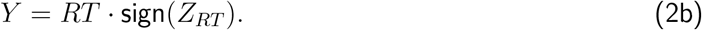

Here, *RT* (2a) is the reaction time and *Y* (2b) is a scalar composite variable encoding both the choice and reaction time, where sign(*Y*) ∈ {−1, +1} is the binary choice (lower if −1, upper if +1) and |*Y* | = *RT >* 0 is the continuous reaction time.

These decision outcomes are stochastic. With the same initial parameters, the diffusion process (1) evolves as a stochastic process with a distribution governed by the probability density function:

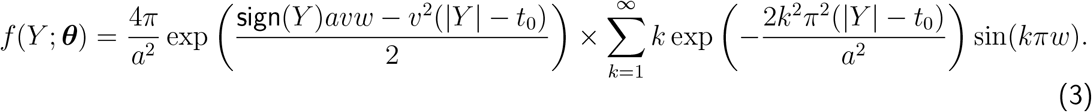

This analytical density was derived in Feller (1968, Chapter 14, Equation 6.15), and later by Navarro & Fuss (2009, Equation 13) who provided the finite approximation of the infinite sum term that we implemented. The relative starting point *z* ∈ (−1, +1) is rewritten as *w* = (1 − sign(*Y*)*z*)*/*2 ∈ (0, +1) to match their notation. For a single trial, the probability of the choice and reaction time pair {sign(*Y*), |*Y* |} is given by this bimodal (one mode per choice; Figure 1C) density function, defined for all |*Y* | *> t*_0_.

#### 2.1.3 Choice and Reaction Time as Functions of Covariates

The DDM parameters ***θ*** = (*a, z, t*_0_, *v*) and consequent decision outcome *Y* may be functions of covariates ***X*** ∈ ℝ^*p*^, such as stimulus properties or experimental conditions (e.g., stimulus strength or reward size):

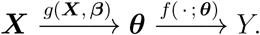

The link functions *g* map covariates to each DDM parameter separately:

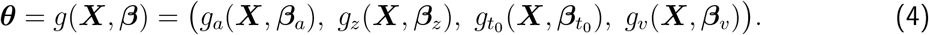

Throughout, ***θ*** refers to *parameters* and ***β*** to *coefficients* to differentiate DDM parameters from covariate effects. For example, take the drift rate to be a linear function 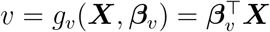: if ***X*** is a scalar *X* = 1, this gives a constant *v* = *β*_*v*,0_; if ***X*** is a vector ***X*** = (1, coherence), then *v* = *β*_*v*,0_ + *β*_*v*,1_coherence, so the drift rate scales with stimulus strength (Figure 1B). The same can apply to *a, z*, and *t*_0_, where the corresponding link functions are similarly defined. Since *a >* 0, *z* ∈ (−1, +1), and *t*_0_ ≥ 0, appropriate link functions (e.g., exponential for *a* and *t*_0_, sigmoid for *z*) map the unbounded linear combination 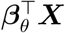 into the valid range of each parameter.

#### 2.1.4 Choices and Reaction Times for a Sequence of Trials

Let ***Y*** = (*Y*_1_, …, *Y*_*n*_) ∈ ℝ^*n*^ denote the sequence of decision outcomes for *n* trials and ***X*** = (***X***_1_, …, ***X***_*n*_) ∈ ℝ^*n×p*^ the covariate matrix, with row ***X***_*i*_ containing the covariates for trial *i* = 1, …, *n*. Applying the covariate model (4) trialwise, the joint density of the sequence can be written as:

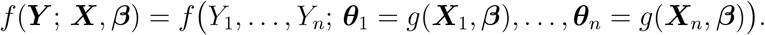

This provides the complete probability specification of ***Y*** and is built from the trialwise marginal densities *f* (*Y*_*i*_; ***θ***_*i*_) defined in (3). The specific factorization depends on whether the marginal densities are treated as independent and/or identically distributed:

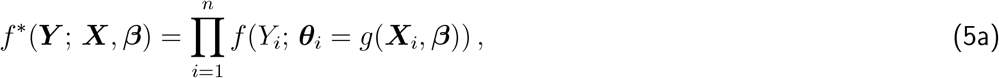

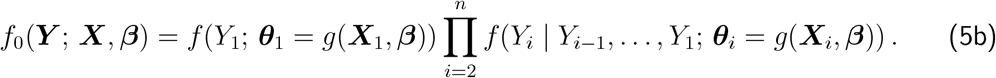

Equation (5a) defines the *independent* case (superscript ∗), in which, conditional on covariates ***X*** and coefficients ***β***, the joint density is the product of trialwise marginal densities. Equation (5b) defines the *dependent* case (subscript 0), in which the joint density factorizes into conditional densities that can depend on the full history of past outcomes. In both cases, non-constant covariates can induce trial-to-trial parameter variability ***θ***_*i*_ = *g*(***X***_*i*_, ***β***) and hence non-identically distributed outcomes. When covariates contribute only a constant intercept *X*_*i*_ = 1 for all *i*, the parameters are fixed ***θ***_*i*_ = ***θ*** and the marginal densities are identical. Then (5a) reduces to the usual *iid* product of identical marginals, while (5b) retains dependence through conditioning on past outcomes.

#### 2.1.5 Misspecification of the Joint Density Function

As discussed, decisions are rarely isolated events; they are richly structured in time with changing internal and external states and influenced by the outcomes of previous decisions. Because of this, it is most realistic to view sequences of trials as neither identically distributed nor independent with joint density function (5b). Going forward, we take this to be the true data-generating density. However, specifying the full data-generating density is often infeasible, as it requires correctly specifying each function that maps measured covariates to DDM parameters, and the complex history effects that induce serial dependence between sequential choices and reaction times.

Instead, for pragmatic reasons, we take the *independent* density (5a) (superscript ∗) as a working approximation of the true density (5b) (subscript 0), accepting the misspecification *f* ^*∗*^ ≠ *f*_0_. By treating trials as independent when they may not be, the working joint density simplifies to the product of trialwise densities conditional only on covariates, rather than on past outcomes. Under this simplification, misspecification can arise from (i) unmodeled DDM parameter variability, (ii) modeled parameter variability with the incorrect functional form (e.g., linear when the true relationship is nonlinear), or (iii) unmodeled serial dependence between trials.

This mismatch between the true and working densities influences estimation and inference, such that the targets of inference are not necessarily the “true” coefficients ***β***_0_ and associated trialwise DDM parameter sequence (***θ***_0,1_, …, ***θ***_0,*n*_) of the complex joint density, but rather the “pseudo-true” coefficients ***β***^*∗*^ and parameters 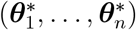 under the independent working density approximation. This best approximation has an intuitive interpretation, in that the “pseudo-true” coefficients are those that minimize the divergence between joint densities *f* ^*∗*^ (5a) and *f*_0_ (5b). Acknowledging that these densities will differ in practice, we formalize the pseudo- (or quasi-) maximum likelihood estimator and explicitly show conditions under which it is consistent and asymptotically normal with a covariance matrix that accounts for this joint-density misspecification. This provides a principled way to quantify estimation uncertainties (e.g., standard errors and confidence intervals) of the “pseudo-true” coefficients under the explicitly general assumptions of unmodeled parameter variability, incorrect functional forms, and/or dependence between trials.

### 2.2 Definitions and Estimation

#### 2.2.1 (Pseudo-) Log-Likelihood Function

The working joint density *f* ^*∗*^(***Y*** ; ***X, β***) (5a) is a function of the decision outcomes ***Y*** given fixed covariates ***X*** and coefficients ***β***. Evaluated at the observed outcomes and re-interpreted as a function of ***β*** (with ***X*** and ***Y*** held fixed), this same product of trialwise densities defines the working likelihood, or “pseudo-likelihood” under misspecification. The expected log-likelihood and its sample average analog (denoted by ^) are defined as:

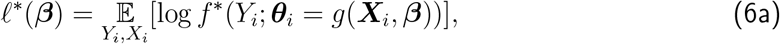

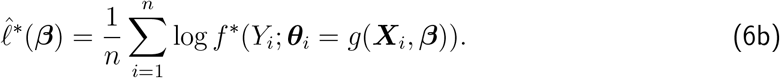

The population expectation (6a) is over the joint distribution of (***X***_*i*_, *Y*_*i*_); the sample analog (6b) conditions on the observed ***X*** as fixed (non-random).

#### 2.2.2 (Pseudo-) Maximum Likelihood Estimator

The “pseudo-true” vector ***β***^*∗*^ contains the coefficients most compatible with the decision outcomes under the possibly misspecified working log-likelihood; equivalently, those that maximize the expected log-likelihood (6a). Similarly, the sample analog 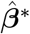 are the coefficients that maximize the sample log-likelihood (6b), i.e., the pseudo-maximum likelihood estimator (MLE):

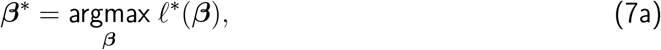

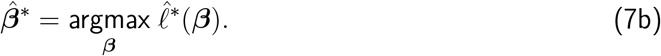

In this paper, we use the *Newton conjugate gradient trust-region algorithm* to iteratively update the estimate of 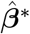 until convergence, when the gradient norm goes below a 10^*−*5^ tolerance. This algorithm moves in the direction of steepest ascent toward the maximum of (6b) using a local quadratic approximation (Steihaug, 1983), as implemented in *SciPy* (Virtanen et al., 2020). This requires first- and second-order derivatives (gradients and Hessians) which are intractable to derive analytically and inefficient to compute numerically with finite differences. Instead, we use automatic differentiation as implemented in *Autograd* (Maclaurin et al., 2015), which propagates derivatives through a computation graph via the chain rule, providing accurate gradients and Hessians that make the log likelihood efficient to optimize.

Numerical issues in estimation can also arise from outliers because extremely fast or slow reaction times have close to zero probability under the DDM density (3), so their log-density terms are close to −∞, destabilizing optimization of the log-likelihood function. A standard remedy is a mixture model that assumes each trial is generated either by the DDM process with probability 1 −*ϵ* or by an outlier process with probability *ϵ*, where the outlier density *f*_outlier_ is taken to be uniform over the interval (min |***Y***|, max |***Y***|) (Ratcliff & Tuerlinckx, 2002). This mixture assigns outliers nonzero probability, preventing extreme log-likelihood penalties and reducing their leverage on the MLE.

### 2.3 Estimator Properties

#### 2.3.1 Assumptions

So far, we have argued that the joint process (***X***_*i*_, *Y*_*i*_) governed by *f*_0_ from (5b) is plausibly neither independent nor identically distributed. However, this level of generality is too broad for large-sample theory, because laws of large numbers and central limit theorems require some form of distributional constancy and restriction on serial dependence. Following Hansen (2022b,a), we adopt the intermediate assumptions of *strict stationarity* and *strong mixing*.

*Strict stationarity* generalizes “identically distributed.” A process is strictly stationary if the joint density of any subset of *k* trials *f*_0_ (***X***_*i*_, *Y*_*i*_), …, (***X***_*i*+*k*_, *Y*_*i*+*k*_) is invariant to the trial index *i* (Hansen, 2022a, Definition 14.2). *Strong mixing* (*α*-mixing) generalizes “independence.” It requires that dependence between events separated by a lag *m* must vanish asymptotically as *m*→ ∞, such that the mixing coefficients which measure this dependence *α*(*m*) →0 (Hansen, 2022a, Section 14.12). Mixing also implies the weaker condition of *ergodicity* (Hansen, 2022a, Theorem 14.14), and the ergodic theorem ensures consistent estimation as a generalization of the weak law of large numbers (Hansen, 2022a, Theorem 14.9). Additionally, under strong mixing with sufficiently fast decay and the required moment conditions, the central limit theorem for correlated observations ensures asymptotic normality of the pseudo-MLE (Hansen, 2022a, Theorem 14.15).

In context, these stability properties are assumptions on the *joint* process (***X***_*i*_, *Y*_*i*_), not on *Y*_*i*_ alone. This distinction is important. The decision outcomes *Y*_*i*_, and the trialwise parameters ***θ***_*i*_ = *g*(***X***_*i*_, ***β***), may be nonstationary, exhibiting trends or drifts across trials. Such nonstationarity does not break the theory, provided it is explicitly modeled by the covariates ***X***_*i*_. For example, if the drift rate *v*_*i*_ trends over time, including the trial number as a covariate (e.g., ***X***_*i*_ = (1, *i*)) allows *g*(***X***_*i*_, ***β***) to model this trend, rendering the joint process (***X***_*i*_, *Y*_*i*_) stationary even if *Y*_*i*_ alone is not. Stationarity and mixing therefore apply to the unexplained variability in *Y*_*i*_ after conditioning on ***X***_*i*_, not to the marginal behavior of *Y*_*i*_ alone. Both properties are preserved under the transformation ***θ***_*i*_ = *g*(***X***_*i*_, ***β***) (Hansen, 2022a, Theorems 14.2,14.12), ensuring the sequence of DDM parameters is also stationary, mixing, and ergodic.

#### 2.3.2 Consistency

The pseudo-MLE 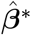 (7b) is the maximizer of the sample log-likelihood (6b), which is itself a sample average version of the expected log-likelihood (6a). Assuming that the underlying process (***X***_*i*_, *Y*_*i*_) is stationary and ergodic (as established above), the log-likelihood contributions log *f* ^*∗*^(*Y*_*i*_; ***θ***_*i*_) are also ergodic transformations (Hansen, 2022a, Theorem 14.5). Consequently, the ergodic theorem applies (Hansen, 2022a, Theorem 14.9), ensuring that the sample log-likelihood converges to the expected log-likelihood:

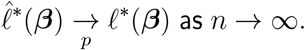

Provided that the “pseudo-true” maximizer ***β***^*∗*^ (7a) is unique (identifiable), such that ^*∗*^(***β***) *<* ^*∗*^(***β***^*∗*^) for all ***β*** ≠ ***β***^*∗*^, the estimator converges to the pseudo-true parameter vector (Hansen, 2022b, Theorem 10.8):

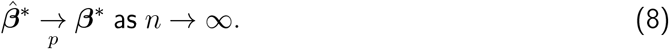

#### 2.3.3 Asymptotic Normality

The uncertainty of the pseudo-MLE is determined by the local curvature of the log-likelihood function at its maximum, where a sharp peak (high curvature) indicates high precision and low variance, while a flat peak (low curvature) indicates greater uncertainty. To formalize its limiting variance, we adapt the central limit theorem for the conventional MLE (Hansen, 2022b, Theorem 10.9) by replacing independent observations with *strict stationarity* and *strong mixing*, and allowing for misspecification of the joint density. Since consistency (8) ensures that 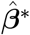 is close to ***β***^*∗*^ for large *n*, we can apply a Taylor approximation of the sample first-order condition 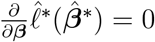 around ***β***^*∗*^, yielding the large-sample approximation:

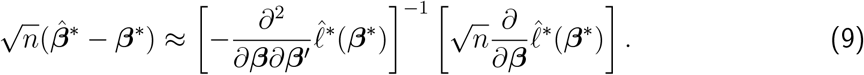

In this expansion, the gradient and Hessian are evaluated at the fixed pseudo-true parameter ***β***^*∗*^, not the random estimator 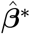, allowing us to apply standard limit theorems to these terms.

The first term in (9) involves the sample Hessian evaluated at ***β***^*∗*^. By the ergodic theorem (Hansen, 2022a, Theorem 14.9), it converges to the *expected Hessian* ℋ_***β***_, defined as the negative expected second derivative:

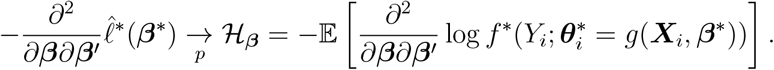

The second term in (9) is the normalized sum of efficient scores. The *efficient score S*_*i*_ is the gradient of the log-likelihood for a single trial, evaluated at ***β***^*∗*^:

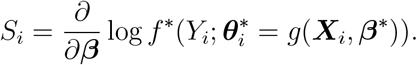

Under *strict stationarity* and *strong mixing*, the score sequence preserves these properties (Hansen, 2022a, Theorems 14.2 and 14.12), and because ***β***^*∗*^ is a maximizer, 𝔼 [*S*_*i*_] = 0. The *Fisher Information Matrix* ℐ_***β***_ is the variance of these scores. While the standard form (without ∼) suffices under independence, serial correlation requires the long-run variance (with ∼), which sums the autocovariances at all lags *k*:

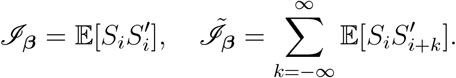

Applying a central limit theorem for correlated observations (Hansen, 2022a, Theorem 14.15), the second term converges to a normal distribution determined by this long-run variance:

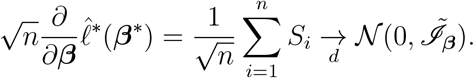

Finally, combining these limits in (9) yields the asymptotic distribution. For correctly specified models, the information matrix equality 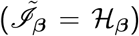 simplifies its covariance. However, under misspecification, this equality does not hold, resulting in a “sandwich” form covariance *V* (Hansen, 2022b, Theorem 10.16):

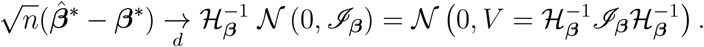

This result accommodates both misspecification, via the failure of the information equality, and serial dependence, via the off-diagonal autocovariance terms in ℐ_***β***_.

#### 2.3.4 Covariance Estimation

In practice, the asymptotic covariance is estimated by replacing population moments with their sample counterparts evaluated at the pseudo-MLE 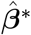. We define the sample Hessian matrix 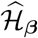, and the sample Fisher Information matrices (short-run 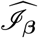 and long-run 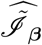):

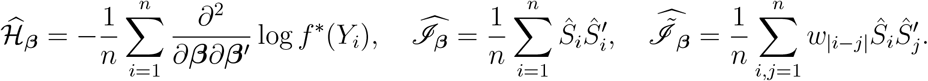

Here, *Ŝ*_*i*_ are the efficient scores evaluated at 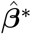, and *w* _|*i* − *j*|_ are kernel weights (e.g., Bartlett) that taper autocovariances as the lag |*i − j*| increases to ensure positive semi-definiteness (Newey & West, 1987).

We consider four estimators of the covariance matrix, ordered from most to least robust:

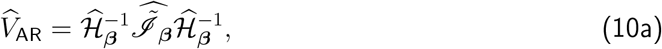

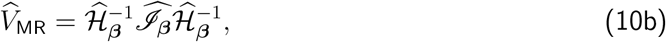

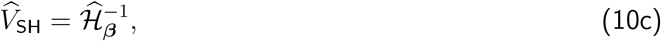

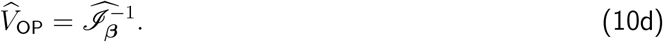

The *Autocorrelation Robust (AR)* estimator (10a) is the most general form, consistent under both misspecification and serial dependence (Newey & West, 1987). The *Misspecification Robust (MR)* estimator (10b), or “sandwich” estimator, allows for misspecification (ℋ_***β***_ /= ℐ_***β***_) but assumes independence 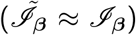 (White, 1982). The *Sample Hessian (SH)* (10c) and *Outer Product (OP)* (10d) estimators rely on the information matrix equality 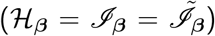, which holds only when the model is correctly specified and the observations are independent (Hansen, 2022b, Theorem 10.5). Standard errors for 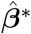 are constructed by taking the square roots of the diagonal elements of 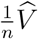.

## 3 Validation by Simulation

### 3.1 Simulation Settings

The finite-sample performance of the pseudo-MLE 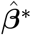 (7b), equivalently 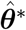 without covariates, and the covariance estimators 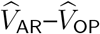 (10a)–(10d) was assessed via Monte Carlo simulations across three settings. In each setting, *b* = 900 independent replications of a sequence of choice and reaction times ***Y*** = (*Y*_1_, …, *Y*_*n*_) of length *n* = 1000 were simulated under the true density (*f*_0_; 5b) and fit under the working density (*f* ^*∗*^; 5a). Settings vary along two dimensions: specification (*f*_0_ = *f* ^*∗*^ or *f*_0_ ≠ *f* ^*∗*^) and parameter structure (fixed or varying across trials), which together determine whether the information matrix equality ℋ_***β***_ = ℐ_***β***_ holds and which covariance estimators are consistent (Section 2.3.4). The first two settings are correctly specified, the second additionally demonstrates modeling of nonstationarity through covariates, while the third provides a realistic test of robustness when parameter variability is present but unmodeled.

- *Correct specification, constant parameters* (*f*_0_ = *f* ^*∗*^): a baseline correctly specified setting with parameters fixed across trials (***θ***_*i*_ = ***θ***^*∗*^, no covariates). Decision outcomes are *iid*, the information matrix equality holds, and all four covariance estimators are expected to be consistent.
- *Correct specification, trending drift rate* (*f*_0_ = *f* ^*∗*^): a correctly specified setting where the drift rate depends linearly on motion coherence via *v*_*i*_ = *g*_*v*_(*X*_*i*_, *β*_*v*_) = *β*_*v*_coherence_*i*_, with coherence decreasing monotonically with trial *i*, while all other parameters are held constant. The information matrix equality holds, and all four estimators are expected to be consistent. This setting demonstrates two capabilities of the pseudo-MLE: modeling DDM parameters as functions of covariates (Section 2.1.3), and modeling nonstationary parameter trends with covariates so that the joint process (*X*_*i*_, *Y*_*i*_) remains stationary (Section 2.3.1).
- *Misspecification, unmodeled parameter variability* (*f*_0_ ≠ *f* ^*∗*^): a misspecified setting where parameters *t*_0_, *v, z* vary *iid* across trials, but the working model assumes constant parameters. This constitutes misspecification of type (i; Section 2.1.5), where the information matrix equality fails and only the sandwich-form estimators 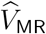 and 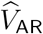 are expected to remain consistent for the covariance around the pseudo-true parameter ***θ***^*∗*^.

All three settings use realistic parameters taken from Matzke & Wagenmakers (2009) (Table 3 Means), rescaled to the present implementation: *ā* = 0.63, 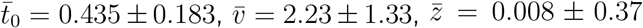, where 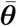 denotes the means of the generating distributions and ± shows variability parameters for *t*_0_, *z* (uniform) and *v* (normal) under misspecification. Under correct specification, these mean values serve directly as the true parameters ***θ***_0_. Under misspecification, however, the pseudo-true parameters ***θ***^*∗*^ are the minimizers of the divergence between the true variable-parameter density and the working constant-parameter density, and therefore need not coincide with the means 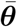 of the generating distributions. Accordingly, ***θ***^*∗*^ for the misspecification setting were determined empirically by fitting the working model to a single large sequence of *n* = 10,000 trials; the smaller *b* = 900 replications of *n* = 1000 trials were then evaluated relative to this reference. Empirical standard deviations and pairwise correlations of 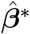 across replications served as ground-truth standard errors and correlations for evaluating the covariance estimators.

### 3.2 Simulation Results

#### 3.2.1 Correct specification, constant parameters

Simulations with constant parameters (Figure 2A,B) serve as a baseline for correct specification (*f*_0_ = *f* ^*∗*^). Point estimates (A; diagonals) are unbiased, meaning the sampling distributions are centered on the true parameters. Pairwise correlations (A; off-diagonals) are positive for (*a, v*) and (*t*_0_, *z*), negative for (*a, t*_0_), (*t*_0_, *v*), and (*v, z*), and near zero for (*a, z*). Analytic standard error estimates (B; diagonals) are mostly unbiased except the non-robust estimators 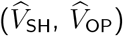 show a slight upward bias for *v*, and the robust estimators 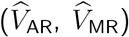 a slight downward bias for *t*_0_ and *z*. The robust estimators also exhibit higher sampling variance than the non-robust estimators, most notably relative to 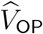, which has the lowest variance overall. Analytic pairwise correlation estimates (B; off-diagonals) are generally unbiased, with slight downward biases for (*a, v*), (*t*_0_, *z*), and (*v, z*) across both robust and non-robust estimators.

**Figure 2:**
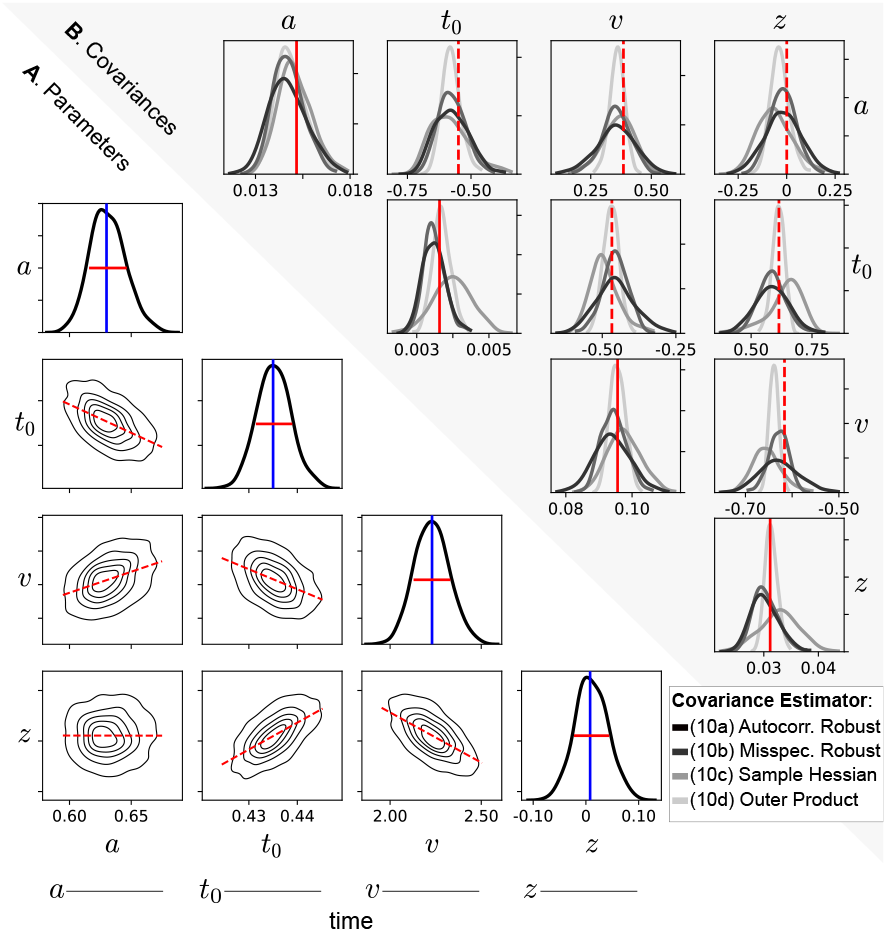
Correct specification, constant parameters. (A) *Diagonals* show sampling distributions of *b* = 900 point estimates (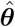; black density curves) around the true parameters (***θ***; blue vertical lines), where ± empirical standard deviations (red solid horizontal lines) are taken to be the true standard errors of the estimator. *Off-diagonals* show bivariate sampling distributions of the point estimates (black density contours), where empirical correlations (red dashed lines) are taken to be the true correlations of the estimator. The schematic below each column indicates how the parameter varies over time (i.e., over the *n* = 1000 trials), in this case indicating that all parameters are constant. (B) *Diagonals* show sampling distributions of the *b* = 900 analytic standard error estimates (i.e., square roots of the diagonal covariance elements; 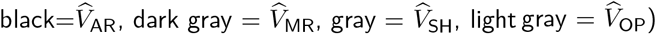) and the empirical standard errors from (A; solid red vertical lines). *Off-diagonals* show analytic pairwise correlation estimates (black-gray density curves) and the empirical correlations from (A; red dashed vertical lines).

#### 3.2.2 Correct specification, trending drift rate

Simulations where the drift rate *v*_*i*_ = *β*_*v*_ ·coherence_*i*_ decreases monotonically with trial index (Figure 3A,B) are also correctly specified (*f*_0_ = *f* ^*∗*^) because nonstationarity is explicitly modeled by the coherence covariate. Results are mostly comparable to the constant parameter simulation above (Figure 2), with the following differences. Point estimates (A; diagonals) remain unbiased, now replacing the parameter *v* with the coefficient *β*_*v*_. Pairwise correlations (A; off-diagonals) are unchanged. Analytic standard error estimates (B; diagonals) are no longer biased upward for *β*_*v*_ as the non-robust estimators were for *v*; instead, 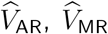, and 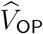 all exhibit a slight downward bias for *t*_0_ and *z*. Analytic pairwise correlation estimates (B; off-diagonals) show fewer biases, with modest downward biases for (*a, t*_0_), (*t*_0_, *z*). These results suggest that the covariance estimators remain mostly accurate when nonstationary trends are explicitly modeled.

**Figure 3:**
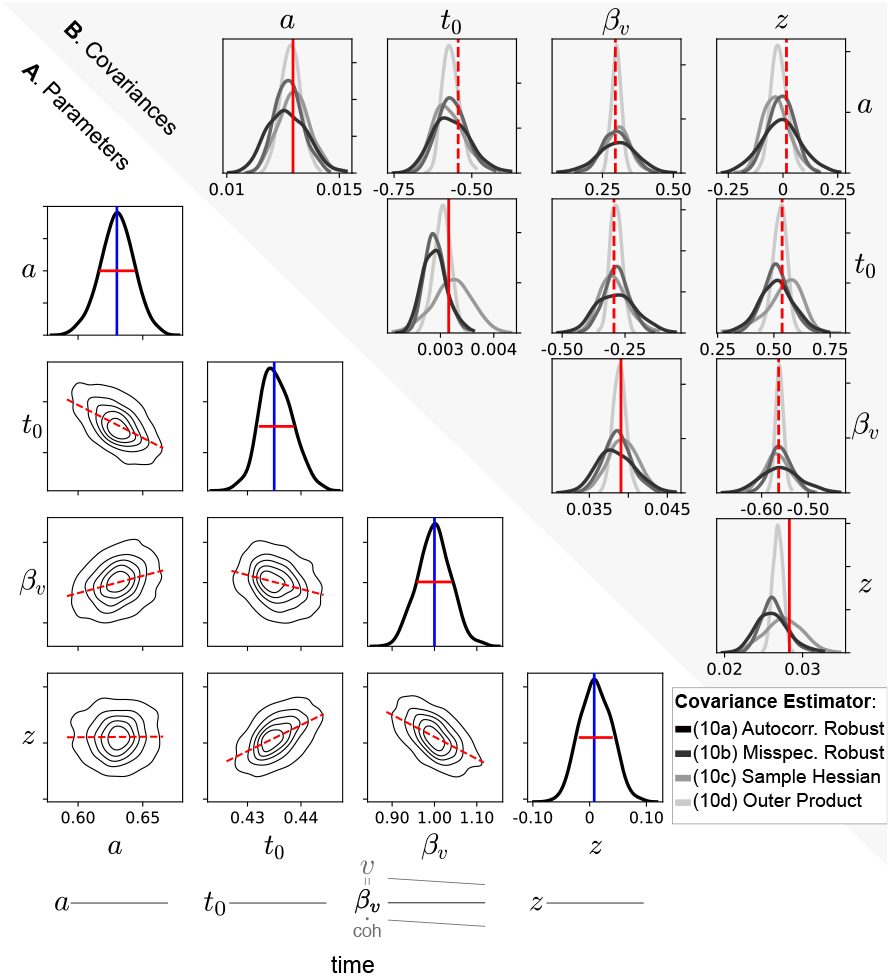
Correct specification, trending drift rate. (A) As in Figure 2A, except the drift rate *v*_*i*_ = *β*_*v*_· coherence_*i*_ decreases monotonically with trial index *i* as coherence declines (i.e., non-stationary), while the coefficient *β*_*v*_ and all other parameters *a, t*_0_, *z* remain constant over time, such that the point estimates are for 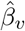 instead of 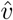; as indicated by the schematic below the third column. (B) As in Figure 2B, except that *β*_*v*_ replaces *v*.

#### 3.2.3 Misspecification, unmodeled parameter variability

Simulations under the seven-parameter DDM, in which *t*_0_, *v*, and *z* are drawn *iid* from distributions centered on 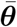 (Figure 4A,B), are misspecified (*f*_0_ ≠ *f* ^*∗*^) because this trial-to-trial variability is unaccounted for in the working model. This is an example of overdispersion, where there is greater variability in the decision outcomes than would be expected under the working model. Point estimates (A; diagonals) show upward bias for *t*_0_ and *z*, and slight downward bias for *a* and *v*, relative to the pseudo-true parameters ***θ***^*∗*^, which do not coincide with 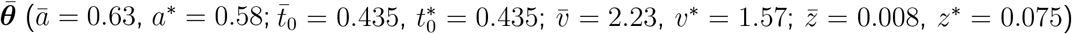. This discrepancy is most apparent for *v* and *z*, which are negatively correlated even under correct specification (Figure 2A), such that the downward shift 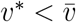 induced by unmodeled variability is compensated by an upward shift 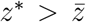. Pairwise correlations (A; off-diagonals) are mostly unchanged, except the positive association for (*a, v*) disappears. Analytic standard error estimates (B; diagonals) show divergence between non-robust and robust estimators, where 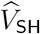 and 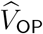 exhibit large downward biases for *t*_0_, *v*, and *z*, which are mitigated, but not fully corrected, by the robust estimators 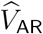 and 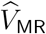. The robust estimators still show a slight downward bias for *z*. Analytic pairwise correlation estimates (B; off-diagonals) follow the same pattern, where non-robust estimators are biased downward for (*a, t*_0_), (*a, z*), (*t*_0_, *z*) and upward for (*a, v*), (*z, v*), with robust estimators largely correcting these biases except for a persistent upward bias in (*z, v*).

**Figure 4:**
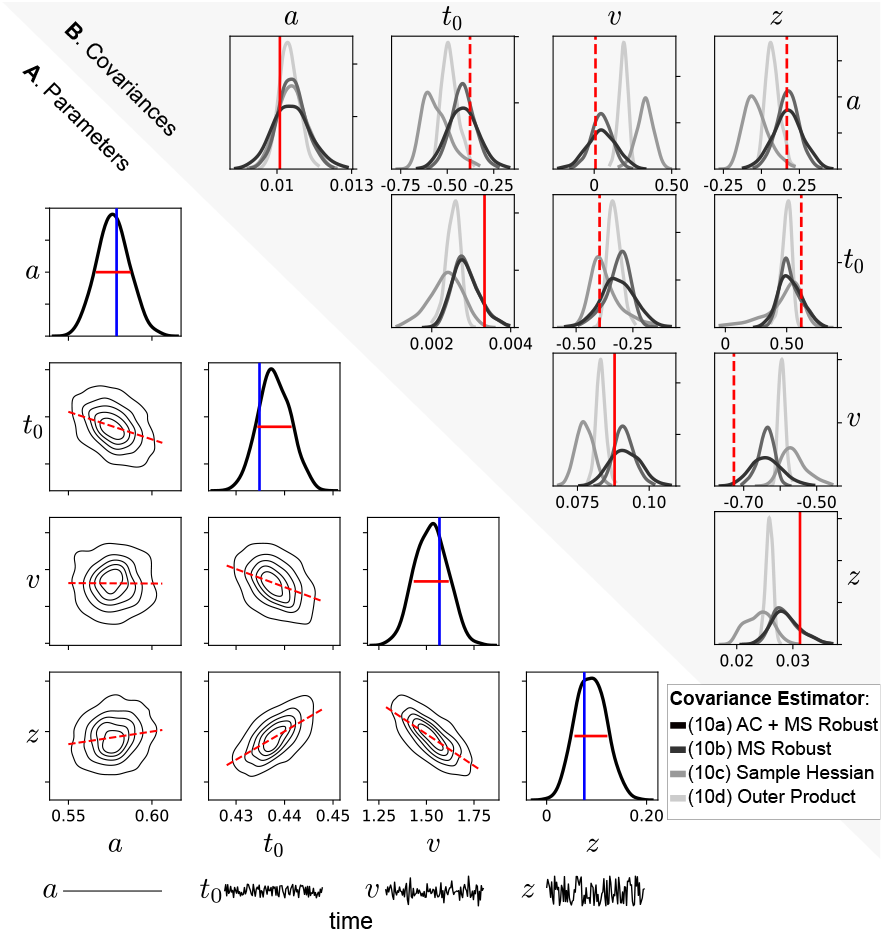
Misspecification, unmodeled parameter variability. (A) As in Figure 2A, except the blue vertical lines now indicate pseudo-true parameters under misspecification. In this simulation, the parameter *a* is constant across time while *t*_0_, *v*, and *z* vary *iid* across time, as indicated by the schematic below each column. (B) As in Figure 2B, except that under this condition, only the 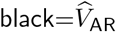 and dark 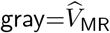 are expected to be consistent estimators under misspecification (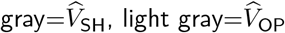 are not).

Overall, across the three simulations, the proposed pseudo-MLE with robust covariances recovered both point estimates and standard errors with minimal bias, implying that the resulting confidence intervals achieve approximately nominal coverage under realistic temporal dependence and parameter variability. This provides a theoretically justified and empirically validated way to fit DDM parameters for which point estimates and confidence intervals remain trustworthy even in the face of unmodeled variability and misspecification, making it possible to distinguish genuine temporal structure in decision states from estimation noise. Such calibrated uncertainty quantification under realistic temporal structure is not available from standard DDM fitting approaches that assume independent, stationary trials, highlighting a capability unique to the proposed method.

## 4 Application to Rat Decision Making

### 4.1 Dataset Description

To illustrate application to empirical data, we applied this method to behavioral data from a self-paced, two-alternative forced-choice (2AFC) reaction time task performed by rats, as detailed in Reinagel (2013). In this experimental paradigm, rats had continuous, 24-hour access to the task from their home cages. Trials were self-initiated via a central start port, triggering a visual stimulus of 100 white dots. A subset of these dots moved coherently toward the left or right (signal), while the remainder moved randomly (noise). Rats viewed the stimulus freely until licking a left or right response port to register their choice, which immediately terminated the stimulus. Correct responses yielded a water reward while error responses resulted in a very small reward.

To isolate intrinsic temporal variability of decision-making parameters from experimentally-induced variability, a contiguous sequence of *n* =42,840 trials was selected from a single rat (ID 195) over a 100-day period, starting after that rat had reached the criterion of 80% correct performance on the task. This sequence was chosen because it demonstrates the nonstationarities and temporal correlations that the proposed method is designed to address, as illustrated below in Figure 5, and is publicly available on Dryad (Reinagel, 2026). Throughout this sequence, task difficulty was held constant at 85% motion coherence, with the direction of motion chosen randomly to be leftward or rightward on each trial.^1^ Consequently, most observed fluctuations in decision behavior are attributable to the passage of time and internal state changes rather than external manipulations. For the *i*^*th*^ trial, stimulus coherence is encoded as the signed covariate *X*_*i*_ ∈ {−0.85, +0.85} (negative for leftward, positive for rightward motion), and the decision outcome as the signed reaction time *Y*_*i*_ (negative for leftward, positive for rightward choice). Elsewhere, the drift rate is modeled as a linear function of coherence *v*_*i*_ = *β*_*v*_*X*_*i*_ (Figure 3), so for consistency, *β*_*v*_ estimates are reported instead of *v* throughout. With the experimental conditions held constant, we looked for evidence of DDM parameters varying systematically over time, across both within-day and between-day timescales.

**Figure 5:**
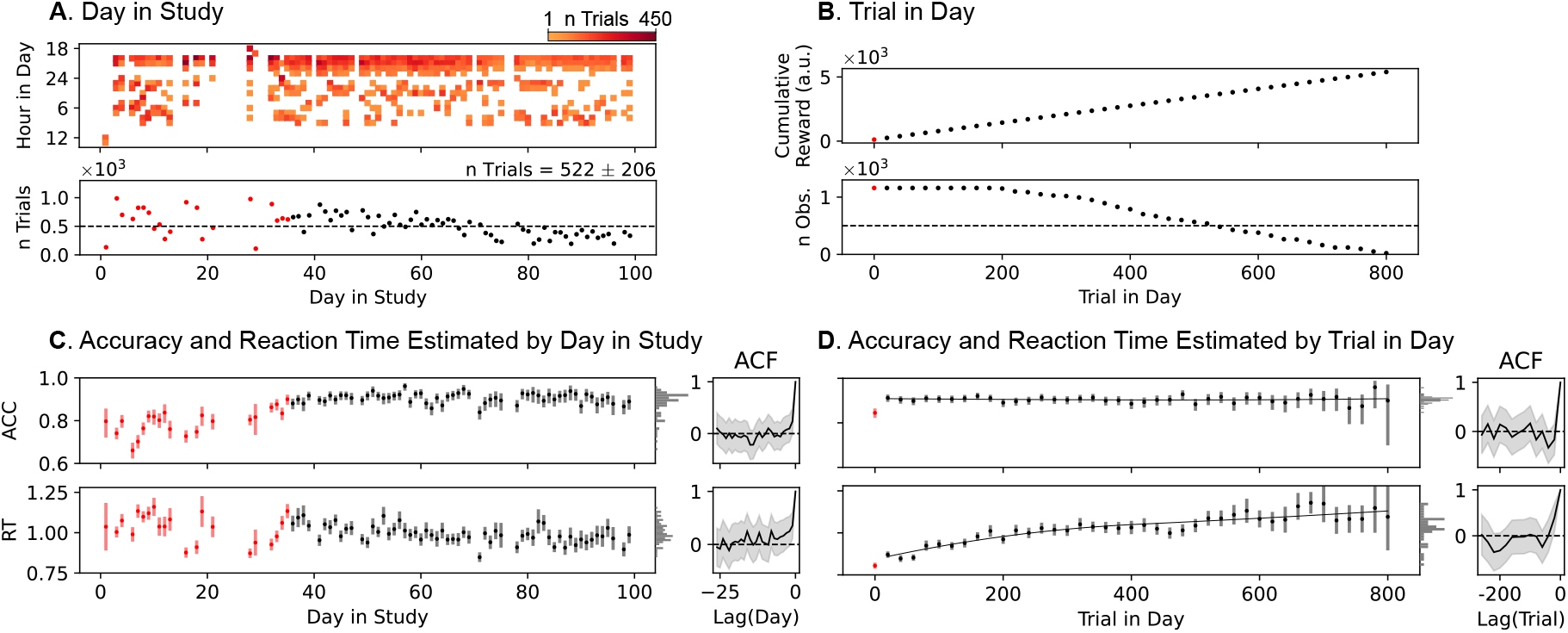
Choices and Reaction Times Reveal Nonstationary Decision Behavior. Behavioral results are displayed for *n*=42,840 random dot motion discrimination trials performed by a single rat over 100 days, beginning after the rat had reached the criterion of 80% correct performance on the task. Left column (A, C) shows between-day temporal structure; right column (B, D) shows within-day temporal structure. A “day” is defined as the 24-hour period beginning at 18:00, which was around the time of day this rat usually chose to start trials. (A; top) Trial heat map showing trial counts per hour (colormap). (A; bottom) Points show trial counts per day (*n* = 491 ± 172). Points in red in A and C (the first 35 days) indicate a period in which the rat continued to improve with practice, before stabilizing thereafter. (B; top) Non-overlapping bins of 20 trials within a day were pooled across days. Points indicate the cumulative water reward received per trial bin, averaged over days. (B; bottom) Points show number of observations (trials) contributing to each point estimate in Panel D. Points in red in B and D (the first 20 trials of each day) are distinguished because accuracy was notably lower in this bin. (C,D) Estimated accuracies and reaction times, with error bars denoting 95% confidence intervals assuming *iid* trials, and either a binomial sampling distribution (for accuracy) or a normal approximation to the sampling distribution (for RT). Insets to the right display autocorrelation functions (ACFs) with 95% pointwise confidence intervals, assuming stationarity. The practice period (red points in C) and first 20 trials of the day (red point in D) were excluded from the ACFs, as they appear nonstationary. Within-day trends in (D) were modeled by fitting piecewise quadratic splines (curves) and then the ACFs were computed on the residuals.

### 4.2 Descriptive Data Analysis

#### 4.2.1 Choices and Reaction Times Reveal Nonstationary Decision Behavior

*Between-day* temporal structure shows this rat typically started performing trials after 18:00, continued vigorously for a few hours, and then performed additional bouts over the course of the night (Figure 5A). Vertical breaks in the heatmap were self-initiated by the rat, while horizontal breaks were experimentally initiated, i.e., the rat was temporarily removed from the task. Over the 100 days, this rat gradually performed fewer and fewer trials per day, a trend explained only by calendar day and no other measured covariates that we checked. Accuracies fluctuated during the first 35 days, showing a gradual increase before plateauing, indicative of improvement with practice; likewise, reaction times were highly variable during these days. From day 36 onward, accuracies continued to fluctuate day-to-day, but less so, and reaction times gradually decreased (Figure 5C). Nonzero autocorrelations persisted for many days, suggesting day-to-day dependencies in choice and reaction times.

*Within-day* cumulative reward increases in proportion to the number of trials performed so far (Figure 5B, upper); therefore, effects of trial-in-day could be explained in part by reward satiety. The rat performed at least 100 trials on every day tested, exceeded 500 trials on half the days, and rarely completed 800 trials in a day. Therefore, the sampling (number of observed trials pooled across days) declined with trial-in-day (Figure 5B, lower). In the first 20 trials of each day, the rat was less accurate and responded more quickly. We speculate that this could be transiently unregulated behavior due to intense thirst, alleviated after a few drops of water were earned. Thereafter, accuracies stabilized but reaction times continued to increase (Figure 5D). Autocorrelations decayed to zero quickly, indicating that fitting the trends with splines accounted for much of the trial-to-trial dependence.

Together these results illustrate that naturalistic decision behavior can contain nonstationarity and temporal fluctuations that are typically present but not modeled. This interesting temporal structure further motivates the estimation of DDM parameters at these timescales. Because the DDM parameters are themselves likely changing on these same timescales, standard inference methods that assume fixed parameters or independent trials may yield biased estimates and overconfident uncertainties. Accounting for this nonstationarity and temporal dependence is therefore not merely a refinement, but a prerequisite for valid inference.

#### 4.2.2 DDM Parameters Reveal Nonstationary Decision States

*Between-day* temporal structure in DDM parameters estimated by day paralleled the behavior, in which parameters fluctuated widely during the initial practice period before stabilizing (Figure 6A, C). In principle, the observed changes in reaction times (Figure 5C) could reflect changes in *a, β*_*v*_, *z*, or some combination thereof. Fitting the DDM by day reveals that, in this case, they are explained by both increases in the decision boundary (*a*) and drift rate coefficient (*β*_*v*_), suggesting the rat became both more cautious and attentive to the motion stimulus with practice. The rat also had a persistent rightward bias (positive *z*) that was strongest at the start of the study and decreased thereafter. All panels show estimates from the proposed pseudo-MLE with confidence intervals (10a) robust to misspecification and temporal dependence, as validated in Section 3. Comparing Figure 6 Panels (A) and (C), the latter controlling for within-day nonstationarity by including trial-in-day spline covariates and reporting each day’s fitted value at the reference trial 400, yielded broadly similar between-day point estimates. Nevertheless, confidence intervals in (C) were often narrower than in (A); across parameters, standard errors were on average 16% smaller. The simulations in Section 3 demonstrate accurate recovery of point estimates and standard errors under realistic temporal dependence and parameter variability, establishing nominal coverage. The narrower intervals here therefore reflect a gain in precision rather than overconfidence, indicating that accounting for within-day nonstationarity mainly improved estimation precision without notably altering the inferred between-day fluctuations. Lastly, nonzero autocorrelations persisted for several days, suggesting day-to-day dependencies in decision states.

**Figure 6:**
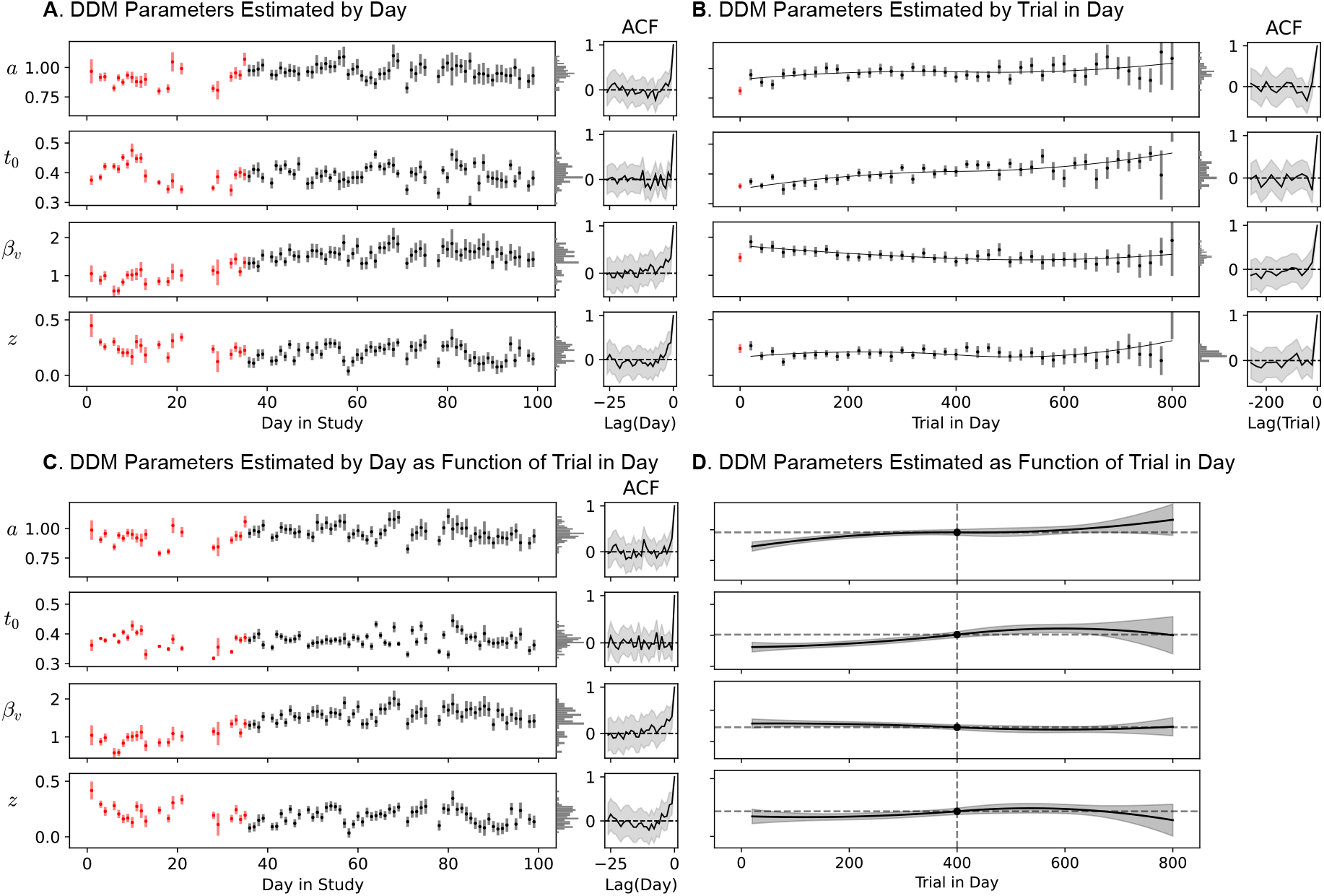
DDM Parameters Reveal Nonstationary Decision States. Estimated DDM parameters and coefficient *β*_*v*_ of motion coherence, corresponding to Figure 5. All panels show pseudo-MLE point estimates with the proposed 95% confidence intervals (10a) robust to temporal dependence and parameter variability, where fluctuations exceeding these intervals indicate non-constant parameters after accounting for estimation uncertainty. Left column (A, C) shows between-day temporal structure; right column (B, D) shows within-day temporal structure. (A) Parameter/coefficient estimates per day in the study; red marks the practice period days and insets show between-day ACFs excluding the points in red. (B) Parameter/coefficient estimates per trial within a day and piecewise quadratic splines fit to point estimates; red marks the first 20 trials of the day and insets show within-day ACFs of the fitted residuals excluding the points in red. However, neither (A) nor (B) explicitly accounts for the within-day nonstationarity visible as trends over trial-in-day. (C, D) Extended analyses addressing this within-day nonstationarity. (D) A single model is fit to all trials (excluding both the first 35 days and first 20 trials of each day, shown in red) where parameters are modeled as piecewise quadratic splines of the trial-in-day covariate. Simultaneous 95% confidence bands are obtained via the delta method applied to (10a), with a Bonferroni correction over the number of unique trial bins per day. The vertical dashed lines show the reference trial 400, and the dashed horizontal lines indicate the corresponding reference value held constant across trials. (C) Extends the between-day analysis of (A) by holding all spline shape coefficients fixed at the values estimated in (D), refitting only the intercept per day, and reporting the fitted value at the reference trial 400, thereby isolating between-day shifts in each DDM parameter while controlling for within-day nonstationarity.

*Within-day* temporal structure in DDM parameters estimated by trial-in-day likewise paralleled the behavior, where the low accuracy and fast reaction times in the first 20 trials of each day (Figure 5D) were explained by a low decision boundary (*a*) and drift rate coefficient (*β*_*v*_; Figure 6B, D). The monotonic increase in reaction times throughout the day (Figure 5D, lower) was explained by both increases in the decision boundary (*a*) and non-decision time (*t*_0_), most apparent in (Figure 6D). This suggests the rat became more cautious and less eager as the day progressed as its need for water was gradually satiated. While the drift rate coefficient (*β*_*v*_) and relative starting point (*z*) in (D) appeared to oscillate, these oscillations were not distinguishable from a constant at this confidence level, highlighting the importance of uncertainty quantification for avoiding misinterpretation of parameter fluctuations. Collectively, the observed trends in DDM parameters reflected changes in latent decision states, especially the monotonically increasing decision bound and non-decision time within a day. These changes are large relative to their uncertainty, indicating that the observed changes in decision states reflect nonstationarity rather than estimation noise.

## 5 Practical Considerations

To contextualize the computational cost and parameter recovery of the proposed pseudo-MLE, we compare it to a Bayesian Markov Chain Monte Carlo (MCMC) estimator from the *HSSM* software (Fengler et al., 2026), the successor to the widely used *HDDM* software (Wiecki et al., 2013). Both methods were applied to the same simulated, correctly specified sequence of choices and reaction times with parameters ***θ*** = {*a* = 0.63, *t*_0_ = 0.435, *v* = 2.23, *z* = 0.008} (Matzke & Wagenmakers, 2009), *n* = {500, 1000, 5000, 10000} trials, and *b* = 100 replications. Likewise, both methods were initialized at the same starting parameter values and constrained to a single CPU core per replication while parallelizing across replications on an 11-core macOS machine; otherwise, default settings were used. *HSSM* (v0.3.0) defaults to 1000 tuning iterations, 1000 posterior draws, and 2 independent chains run sequentially. We compare runtime in seconds (averaged across replications), and root mean squared error (RMSE) of parameter and standard error estimates:

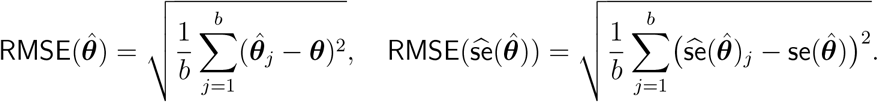

Here, 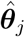 is the point estimate for the *j*-th replication, 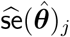 is its estimated standard error (standard error for MLE, marginal posterior standard deviation for MCMC), and 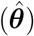 is the empirical standard deviation of the *b* parameter estimates, serving as the true standard error.

Theoretically, parameters and standard errors are different when comparing MLE (Frequentist) and MCMC (Bayesian) methods, as the latter defines a prior on the parameter space. Additionally, MCMC estimates the full posterior distribution from which point estimates (marginal posterior means) and their standard errors (marginal posterior standard deviations) are calculated. Yet the influence of the prior diminishes at large samples, and asymptotically, the posterior converges to a normal distribution centered on the MLE with covariance 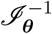 (van der Vaart, 1998, Theorem 10.1); the same as the MLE under *iid* trials and correct specification in Section 2.3. This theorem links MLE and MCMC inference at large samples, and therefore we can compare the RMSE of parameter and standard error estimates of these methods.

Figure 7 compares MLE and MCMC under *iid* trials and correct specification, across four sequence lengths. MCMC takes 9–43× longer, with the gap widening at larger *n* as the per-sample likelihood evaluation cost compounds over many posterior samples. MLE is more computationally efficient because it uses first- and second-order likelihood information to converge directly to a point estimate, whereas MCMC must characterize the full posterior distribution through stochastic sampling. Despite this difference in computational cost, both methods recover parameters and their standard errors accurately, where both improve at larger *n*, and MLE performs as well or better than MCMC across most settings (except for 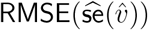 at *n* = 1000 trials).

**Figure 7:**
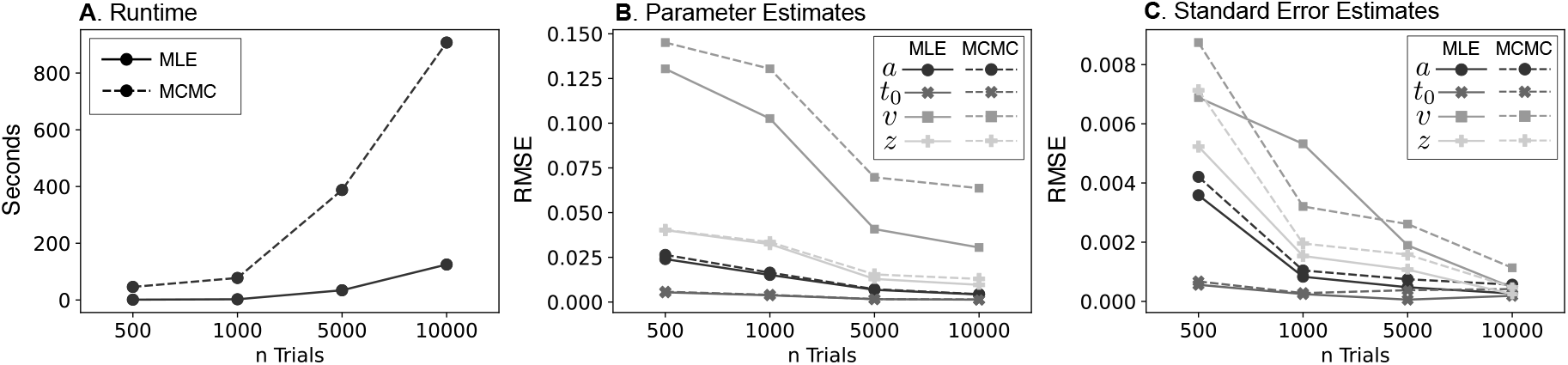
Comparison of MLE and MCMC. Solid lines denote MLE, dashed lines denote MCMC, and different marker shapes distinguish the four parameters *a, t*_0_, *v*, and *z* across all panels. (A) Average runtime in seconds across replications as a function of sample size. (B) RMSE of the parameter estimates, in the original units of *a, t*_0_, *v*, and *z*. (C) RMSE of the standard error estimates.

In context, both methods allow for covariate modeling and uncertainty quantification; however, MCMC as implemented assumes *iid* trials and correct specification (Wiecki et al., 2013; Moens & Zenon, 2018). Therefore, standard errors are likely underestimated under temporal dependence and unmodeled parameter variability, while we show that pseudo-MLE robustly estimates standard errors in this setting (Figure 4). That said, their MCMC approach has a lot to offer that we cannot. Most notably, the *HSSM* software can approximate likelihoods when no analytical form exists, differentiable or not (Fengler et al., 2026, 2022, 2021). This allows for a wide class of DDMs beyond the four-parameter model, including the seven-parameter model (Ratcliff & Rouder, 1998; Ratcliff & Tuerlinckx, 2002), collapsing-bound model (Drugowitsch et al., 2012), Lévy-flight and Cauchy-flight models (Voss et al., 2019). Also, it can implement hierarchical structures to pool data across subjects or sessions, improving estimation accuracy when individual data is sparse. Therefore, while Bayesian MCMC offers extensive flexibility for complex and hierarchical generative models, our pseudo-MLE approach provides a highly scalable, explicitly robust alternative for fast inference and uncertainty quantification under temporal dependence and parameter variability.

## 6 Conclusions

Standard DDM fitting methods treat trial sequences as independent and identically distributed (*iid*), an assumption routinely violated in naturalistic behavioral experiments where decisions are richly structured in time (Urai, 2025; Moens & Zenon, 2018). This paper introduces a pseudo-MLE method that provides analytically tractable uncertainty quantification for DDM parameters while remaining valid under temporal dependence and unmodeled parameter variability, and extending to settings where parameters are explicitly modeled as functions of covariates.

### Theoretical Contributions

We establish that the pseudo-MLE converges to its pseudo-true coefficients and is asymptotically normal under strict stationarity and strong mixing. The key insight is that stationarity applies to the joint process of covariates and outcomes, not to the decision outcomes alone. By explicitly modeling structured nonstationarity — such as session-level trends, circadian rhythms, and learning effects — valid point estimates and uncertainties are recovered even in nonstationary examples, substantially broadening the scope of settings to which the method applies. The resulting sandwich-form covariance estimators (Newey & West, 1987; Hansen, 2022b) correct for the failure of the information matrix equality under possible misspecification, providing valid standard errors and confidence intervals even when the working model is an approximation of the true data-generating process.

### Simulation Results

Monte Carlo simulations confirm that under correct specification, with either constant or covariate-driven nonstationary parameters, point estimates are unbiased and covariances are accurately recovered, consistent with prior DDM parameter recovery studies (Boehm et al., 2018; Matzke & Wagenmakers, 2009). Under misspecification induced by unmodeled trial-to-trial parameter variability, non-robust covariance estimators substantially underestimate standard errors, while robust estimators largely mitigate these biases. Robust covariance estimation is therefore advisable whenever misspecification cannot be ruled out, which Urai (2025) cautions is the norm in decision-making experiments.

More broadly, explicitly modeling parameter variability via covariates is preferable to absorbing it into unstructured noise parameters, which carry no temporally structured information and conflate distinct underlying sources such as arousal, choice history, and learning (Duffy et al., 2025; Urai, 2025; Roy et al., 2018, 2021). A notable example is the seven-parameter DDM, where trial-to-trial variability in boundary height *a* and starting point *z* is difficult to disentangle, potentially leading to misattributed variability (Boehm et al., 2018; Lerche & Voss, 2016; Ratcliff et al., 2018). Covariate modeling offers a more tractable alternative: by linking *a* and *z* separately to observed covariates, their respective sources of variability can be more cleanly separated. More generally, when structured variability can be tied to observable quantities, covariate modeling converts a source of inferential noise into interpretable signal about how decision states change over time.

### Data Analysis Results

Applied to empirical data from a single rat performing a visual motion discrimination task (Reinagel, 2013), the method revealed systematic multi-timescale structure in DDM parameters invisible to standard analyses. At the between-day timescale, increasing decision boundaries and drift rate coefficients across the first 35 days could be a practice effect as often reported (Ashwood et al., 2022; Mohammadi et al., 2025; Cochrane et al., 2023). Notably, parameters continued to vary well beyond this initial period, suggesting that decision states change even in well-trained animals. Comparing Figure 6A,C further shows that explicitly modeling within-day nonstationary trends did not materially change the between-day point estimates, but it did sharpen inference by reducing uncertainty; across parameters, standard errors were smaller when trial-in-day covariates were included. Within a day, decision boundaries and non-decision times increased monotonically as the rat became satiated and fatigued, while markedly lower boundaries and faster responses in the first 20 trials might reflect heightened thirst at the start of the day. These findings show that even under mostly constant experimental conditions, DDM parameters varied substantially relative to their uncertainties, changes that cannot be inferred from point estimates alone and that methods ignoring trial order would attribute to noise. Persistent auto-correlations across trials and days further confirmed that sequential dependencies are present and should be accounted for in inference, consistent with history effects documented across rodent and primate decision making (Urai et al., 2019; Busse et al., 2011; Danskin et al., 2023; Frund et al., 2014; Gao et al., 2009).

### Limitations and Future Directions

While misspecification-robust covariances reduce inferential errors, they cannot replace the explicit modeling of known sources of variability. For instance, history-dependent biases that drive sequential dependencies were not modeled here, as they were in (Nguyen et al., 2019; Urai et al., 2019; Braun et al., 2018; Gupta et al., 2024). Methodologically, the large-sample theory underlying our pseudo-MLE approach requires that the joint process of covariates and outcomes be strictly stationary and strongly mixing. These conditions are difficult to verify in practice, although our application to rat decision making suggests that realistic within-day nonstationarity did not substantively distort the between-day point estimates in this example (Figure 6A,C); explicitly modeling it improved estimation precision. Empirically, applying the method to a single rat under fixed stimulus conditions was meant as a demonstration, and the results presented are specific to this example and not expected to generalize across different rats or task designs. We analyzed example covariates by way of illustration of the method, but did not attempt to exhaust the many potentially explanatory covariates in this dataset. Finally, because the current method estimates fixed coefficients within each fitted window, it characterizes continuous parameter shifts but cannot capture the abrupt, discrete strategy switches that latent-state models are specifically designed to detect (Ashwood et al., 2022; Mohammadi et al., 2025; Kucharský et al., 2021).

Future work could address these limitations in three primary ways. First, history-dependent biases and species-specific traits (Nguyen & Reinagel, 2022; Shevinsky & Reinagel, 2019; Li et al., 2024; Pagan et al., 2025) are better addressed by incorporating them as covariates into the underlying model. Extending the method to large, multi-subject datasets would then enable the comparison of changing decision dynamics across different subjects, timescales, and species (Asadpour et al., 2024; Danskin et al., 2023; Nguyen & Reinagel, 2022). Second, testing additional covariates derived from external behavioral metrics, such as pose tracking (Mathis et al., 2018) or arousal signals (Murphy et al., 2014), would give richer insights into how physical context shapes evidence accumulation. Linking these physical signals directly to DDM parameters would further disentangle structured variability explained by observed covariates and residual variability that requires robust inference. Finally, combining our pseudo-MLE approach with latent-state models would more fully capture the broad spectrum of temporal variability, such as hidden Markov models for discrete state switches (Ashwood et al., 2022; Mohammadi et al., 2025; Kucharský et al., 2021), autoregressive processes for gradual drift (Urai et al., 2019; Vloeberghs et al., 2025), transition models that combine discrete switches with continuous drift (Schumacher et al., 2023, 2025), or reinforcement learning for history effects (Pedersen et al., 2017; Fengler et al., 2022). Together, these extensions could jointly account for structured nonstationarity, gradual parameter drift, and discrete state changes, more fully capturing the broad spectrum of temporal variability inherent in sequential decision making.

In conclusion, this work establishes a methodological foundation for studying dynamic decision making in naturalistic settings, in theory and applications. By relaxing the standard assumptions of correct specification and trial independence, and by grounding stationarity in the joint process of covariates and outcomes rather than in decision outcomes alone, the pseudo-MLE method enables principled and computationally efficient inference about sequential decision making in naturalistic experiments.

## 7 Code and Data Availability

The Python source code for the methods described in this paper is publicly available on GitHub (https://github.com/griegner/drift-diffusion). The data, scripts, and computational environment used to reproduce the results reported in this paper are publicly available on Code Ocean (https://doi.org/10.24433/CO.5868634.v1).

## 8 Acknowledgements

We would like to thank Mikio Aoi (University of California, San Diego) for helpful feedback on the manuscript. This work was supported by the National Institutes of Health (NIH) BRAIN Initiative grant 1R34NS132037-01.

## Footnotes

1 There were two experimentally imposed one-trial-back sequential dependencies: rewards following correct trials were larger than rewards following error trials, and correction trials were imposed after some errors.

## References

Asadpour, A., Tan, H., Lenfesty, B., & Wong-Lin, K. (2024). Of Rodents and Primates: Time-Variant Gain in Drift–Diffusion Decision Models. Computational Brain & Behavior, 7(2), 195–206. 10.1007/s42113-023-00194-1

Ashwood, Z. C., Roy, N. A., Stone, I. R., The International Brain Laboratory, Urai, A. E., Churchland, A. K., Pouget, A., & Pillow, J. W. (2022). Mice alternate between discrete strategies during perceptual decision-making. Nature Neuroscience, 25(2), 201–212. 10.1038/s41593-021-01007-z

Boehm, U., Annis, J., Frank, M. J., Hawkins, G. E., Heathcote, A., Kellen, D., Krypotos, A.-M., Lerche, V., Logan, G. D., Palmeri, T. J., van Ravenzwaaij, D., Servant, M., Singmann, H., Starns, J. J., Voss, A., Wiecki, T. V., Matzke, D., & Wagenmakers, E.-J. (2018). Estimating across-trial variability parameters of the Diffusion Decision Model: Expert advice and recommendations. Journal of Mathematical Psychology, 87, 46–75.

Bogacz, R., Brown, E., Moehlis, J., Holmes, P., & Cohen, J. D. (2006). The physics of optimal decision making: A formal analysis of models of performance in two-alternative forced-choice tasks. Psychological Review, 113(4), 700–765. 10.1037/0033-295X.113.4.700

Braun, A., Urai, A. E., & Donner, T. H. (2018). Adaptive History Biases Result from Confidence-Weighted Accumulation of past Choices. The Journal of Neuroscience, 38(10), 2418–2429. 10.1523/JNEUROSCI.2189-17.2017

Busse, L., Ayaz, A., Dhruv, N. T., Katzner, S., Saleem, A. B., Schölvinck, M. L., Zaharia, A. D., & Carandini, M. (2011). The Detection of Visual Contrast in the Behaving Mouse. The Journal of Neuroscience, 31(31), 11351–11361. 10.1523/JNEUROSCI.6689-10.2011

Cochrane, A., Sims, C. R., Bejjanki, V. R., Green, C. S., & Bavelier, D. (2023). Multiple timescales of learning indicated by changes in evidence-accumulation processes during perceptual decision-making. npj Science of Learning, 8(1), 19. 10.1038/s41539-023-00168-9

Danskin, B. P., Hattori, R., Zhang, Y. E., Babic, Z., Aoi, M., & Komiyama, T. (2023). Exponential history integration with diverse temporal scales in retrosplenial cortex supports hyperbolic behavior. Science Advances, 9(48), eadj4897. 10.1126/sciadv.adj4897

Deakin, J., Schofield, A., & Heinke, D. (2024). Support for the Time-Varying Drift Rate Model of Perceptual Discrimination in Dynamic and Static Noise Using Bayesian Model-Fitting Methodology. Entropy, 26(8), 642. 10.3390/e26080642

Drugowitsch, J., Moreno-Bote, R., Churchland, A. K., Shadlen, M. N., & Pouget, A. (2012). The Cost of Accumulating Evidence in Perceptual Decision Making. The Journal of Neuroscience, 32(11), 3612–3628. 10.1523/JNEUROSCI.4010-11.2012

Duffy, J. S., Bellgrove, M. A., Murphy, P. R., & O’Connell, R. G. (2025). Disentangling sources of variability in decision-making. Nature Reviews Neuroscience, 26(5), 247–262. 10.1038/s41583-025-00916-3

Feller, W. (1968). An introduction to probability theory and its applications (third ed. rev ed.). Wiley series in probability and mathematical statistics. J. Wiley.

Fengler, A., Bera, K., Pedersen, M. L., & Frank, M. J. (2022). Beyond drift diffusion models: fitting a broad class of decision and reinforcement learning models with HDDM. Journal of Cognitive Neuroscience, 34(10), 1780–1805. 10.1162/jocn_a_01902

Fengler, A., Govindarajan, L. N., Chen, T., & Frank, M. J. (2021). Likelihood approximation networks (LANs) for fast inference of simulation models in cognitive neuroscience. eLife, 10, e65074. 10.7554/eLife.65074

Fengler, A., Xu, Y., Bera, K., & Frank, M. (2026). HSSM: A generalized toolbox for hierarchical bayesian estimation of computational models in cognitive neuroscience. In Preparation.

Frund, I., Wichmann, F. A., & Macke, J. H. (2014). Quantifying the effect of intertrial dependence on perceptual decisions. Journal of Vision, 14(7), 9–9. 10.1167/14.7.9

Gao, J., Wong-Lin, K., Holmes, P., Simen, P., & Cohen, J. D. (2009). Sequential Effects in Two-Choice Reaction Time Tasks: Decomposition and Synthesis of Mechanisms. Neural Computation, 21(9), 2407–2436. 10.1162/neco.2009.09-08-866

Gunawan, D., Hawkins, G. E., Kohn, R., Tran, M.-N., & Brown, S. D. (2022). Time-evolving psychological processes over repeated decisions. Psychological Review, 129(3), 438–456. 10.1037/rev0000351

Gupta, D., DePasquale, B., Kopec, C. D., & Brody, C. D. (2024). Trial-history biases in evidence accumulation can give rise to apparent lapses in decision-making. Nature Communications, 15(1), 662. 10.1038/s41467-024-44880-5

Hansen, B. E. (2022a). Econometrics. Princeton University Press.

Hansen, B. E. (2022b). Probability and statistics for economists. Princeton University Press.

Hato, T., Schumacher, L., Radev, S. T., & Voss, A. (2025). Lévy Versus Wiener: Assessing the Effects of Model Misspecification on Diffusion Model Parameters. Computational Brain & Behavior. 10.1007/s42113-025-00248-6

Kucharský, S., Tran, N.-H., Veldkamp, K., Raijmakers, M., & Visser, I. (2021). Hidden Markov Models of Evidence Accumulation in Speeded Decision Tasks. Computational Brain & Behavior, 4(4), 416–441. 10.1007/s42113-021-00115-0

Lerche, V. & Voss, A. (2016). Model Complexity in Diffusion Modeling: Benefits of Making the Model More Parsimonious. Frontiers in Psychology, 7. 10.3389/fpsyg.2016.01324

Lerche, V., Voss, A., & Nagler, M. (2017). How many trials are required for parameter estimation in diffusion modeling? A comparison of different optimization criteria. Behavior Research Methods, 49(2), 513–537. 10.3758/s13428-016-0740-2

Li, J.-J., Shi, C., Li, L., & Collins, A. G. (2024). Dynamic noise estimation: A generalized method for modeling noise fluctuations in decision-making. Journal of Mathematical Psychology, 119, 102842. 10.1016/j.jmp.2024.102842

Maclaurin, D., Duvenaud, D., & Adams, R. P. (2015). Autograd: Effortless Gradients in Numpy. ICML 2015 AutoML Workshop, 238, 5.

Maggi, S., Hock, R. M., O’Neill, M., Buckley, M., Moran, P. M., Bast, T., Sami, M., & Humphries, M. D. (2024). Tracking subjects’ strategies in behavioural choice experiments at trial resolution. eLife, 13, e86491. 10.7554/eLife.86491

Mathis, A., Mamidanna, P., Cury, K. M., Abe, T., Murthy, V. N., Mathis, M. W., & Bethge, M. (2018). DeepLabCut: markerless pose estimation of user-defined body parts with deep learning. Nature Neuroscience, 21(9), 1281–1289. 10.1038/s41593-018-0209-y

Matzke, D. & Wagenmakers, E.-J. (2009). Psychological interpretation of the ex-Gaussian and shifted Wald parameters: A diffusion model analysis. Psychonomic Bulletin & Review, 16(5), 798–817. 10.3758/PBR.16.5.798

Moens, M. & Zenon, A. (2018). Recurrent Auto-Encoding Drift Diffusion Model. bioRxiv. 10.1101/220517

Mohammadi, Z., Ashwood, Z. C., The International Brain Laboratory, & Pillow, J. W. (2025). Identifying the factors governing internal state switches during nonstationary sensory decision-making. Nature Communications, 16(1), 11684. 10.1038/s41467-025-66738-0

Murphy, P. R., Vandekerckhove, J., & Nieuwenhuis, S. (2014). Pupil-Linked Arousal Determines Variability in Perceptual Decision Making. PLoS Computational Biology, 10(9), e1003854. 10.1371/journal.pcbi.1003854

Navarro, D. J. & Fuss, I. G. (2009). Fast and accurate calculations for first-passage times in Wiener diffusion models. Journal of Mathematical Psychology, 53(4), 222–230. 10.1016/j.jmp.2009.02.003

Newey, W. & West, K. (1987). A simple, positive semi-definite, heteroskedasticity and autocor-relationconsistent covariance matrix. Econometrica, 55(3), 703–08.

Nguyen, K. P., Josić, K., & Kilpatrick, Z. P. (2019). Optimizing sequential decisions in the drift–diffusion model. Journal of Mathematical Psychology, 88, 32–47. 10.1016/j.jmp.2018.11.001

Nguyen, Q. N. & Reinagel, P. (2022). Different Forms of Variability Could Explain a Difference Between Human and Rat Decision Making. Frontiers in Neuroscience, 16, 794681. 10.3389/fnins.2022.794681

Pagan, M., Tang, V. D., Aoi, M. C., Pillow, J. W., Mante, V., Sussillo, D., & Brody, C. D. (2025). Individual variability of neural computations underlying flexible decisions. Nature, 639(8054), 421–429. 10.1038/s41586-024-08433-6

Pedersen, M. L., Frank, M. J., & Biele, G. (2017). The drift diffusion model as the choice rule in reinforcement learning. Psychonomic Bulletin & Review, 24(4), 1234–1251. 10.3758/s13423-016-1199-y

Perquin, M. N., Heed, T., & Kayser, C. (2024). Variance (un)explained: Experimental conditions and temporal dependencies explain similarly small proportions of reaction time variability in linear models of perceptual and cognitive tasks. Journal of Experimental Psychology: General, 153(12), 3107–3129. 10.1037/xge0001630

Perquin, M. N., Van Vugt, M. K., Hedge, C., & Bompas, A. (2023). Temporal Structure in Sensorimotor Variability: A Stable Trait, But What For? Computational Brain & Behavior, 6(3), 400–437. 10.1007/s42113-022-00162-1

Ratcliff, R. (1978). A Theory of Memory Retrieval. Psychological Review, 85(2).

Ratcliff, R. (2013). Parameter variability and distributional assumptions in the diffusion model. Psychological Review, 120(1), 281–292. 10.1037/a0030775

Ratcliff, R. & Childers, R. (2015). Individual differences and fitting methods for the two-choice diffusion model of decision making. Decision, 2(4), 237–279. 10.1037/dec0000030

Ratcliff, R. & Rouder, J. N. (1998). Modeling Response Times for Two-Choice Decisions. Psychological Science, 9(5), 347–356. 10.1111/1467-9280.00067

Ratcliff, R. & Tuerlinckx, F. (2002). Estimating parameters of the diffusion model: Approaches to dealing with contaminant reaction times and parameter variability. Psychonomic Bulletin & Review, 9(3), 438–481. 10.3758/BF03196302

Ratcliff, R., Voskuilen, C., & McKoon, G. (2018). Internal and external sources of variability in perceptual decision-making. Psychological Review, 125(1), 33–46. 10.1037/rev0000080

Reinagel, P. (2013). Speed and Accuracy of Visual Motion Discrimination by Rats. PLoS ONE, 8(6), e68505. 10.1371/journal.pone.0068505

Reinagel, P. (2026). Rat visual perceptual decision making: A 100-day longitudinal behavioral time series (in press). 10.5061/dryad.hhmgqnkxb

Roy, N. A., Bak, J. H., Akrami, A., Brody, C. D., & Pillow, J. W. (2018). Efficient inference for time-varying behavior during learning. Advances in Neural Information Processing Systems, 31, 5695–5705.

Roy, N. A., Bak, J. H., Akrami, A., Brody, C. D., & Pillow, J. W. (2021). Extracting the dynamics of behavior in sensory decision-making experiments. Neuron, 109(4), 597–610.e6. 10.1016/j.neuron.2020.12.004

Schumacher, L., Bürkner, P.-C., Voss, A., Köthe, U., & Radev, S. T. (2023). Neural superstatistics for Bayesian estimation of dynamic cognitive models. Scientific Reports, 13(1), 13778. 10.1038/s41598-023-40278-3

Schumacher, L., Schnuerch, M., Voss, A., & Radev, S. T. (2025). Validation and Comparison of Non-stationary Cognitive Models: A Diffusion Model Application. Computational Brain & Behavior, 8(2), 191–210. 10.1007/s42113-024-00218-4

Shadlen, M. N., Churchland, A. K., & Yang, T. (2006). The Speed and Accuracy of a Simple Perceptual Decision: A Mathematical Primer. Bayesian brain: Probabilistic approaches to neural coding, 209–237.

Shadlen, M. N. & Newsome, W. T. (2001). Neural Basis of a Perceptual Decision in the Parietal Cortex (Area LIP) of the Rhesus Monkey. Journal of Neurophysiology, 86(4), 1916–1936. 10.1152/jn.2001.86.4.1916

Shevinsky, C. A. & Reinagel, P. (2019). The Interaction Between Elapsed Time and Decision Accuracy Differs Between Humans and Rats. Frontiers in Neuroscience, 13, 1211. 10.3389/fnins.2019.01211

Shinn, M., Lam, N. H., & Murray, J. D. (2020). A flexible framework for simulating and fitting generalized drift-diffusion models. eLife, 9, e56938. 10.7554/eLife.56938

Steihaug, T. (1983). The Conjugate Gradient Method and Trust Regions in Large Scale Optimization. SIAM Journal on Numerical Analysis, 20(3), 626–637. 10.1137/0720042

Steinemann, N., Stine, G. M., Trautmann, E., Zylberberg, A., Wolpert, D. M., & Shadlen, M. N. (2024). Direct observation of the neural computations underlying a single decision. eLife, 12, RP90859. 10.7554/eLife.90859

Stone, M. (1960). Models for Choice-Reaction Time. Psychometrika, 25(3), 251–260. 10.1007/BF02289729

Treviño, M., Medina-Coss Y León, R., & Lezama, E. (2022). Response Time Distributions and the Accumulation of Visual Evidence in Freely Moving Mice. Neuroscience, 501, 25–41. 10.1016/j.neuroscience.2022.08.015

Urai, A. E. (2025). Structure uncovered: understanding temporal variability in perceptual decision-making. Trends in Cognitive Sciences, S1364661325001494. 10.1016/j.tics.2025.06.003

Urai, A. E., De Gee, J. W., Tsetsos, K., & Donner, T. H. (2019). Choice history biases subsequent evidence accumulation. eLife, 8, e46331. 10.7554/eLife.46331

van der Vaart, A. (1998). Asymptotic Statistics (1 ed.). Cambridge University Press. 10.1017/CBO9780511802256

Van Ravenzwaaij, D. & Oberauer, K. (2009). How to use the diffusion model: Parameter recovery of three methods: EZ, fast-dm, and DMAT. Journal of Mathematical Psychology, 53(6), 463–473. 10.1016/j.jmp.2009.09.004

Virtanen, P., Gommers, R., Oliphant, T. E., Haberland, M., Reddy, T., Cournapeau, D., Burovski, E., Peterson, P., Weckesser, W., & Bright, J. (2020). SciPy 1.0: fundamental algorithms for scientific computing in Python. Nature Methods, 17(3), 261–272. 10.1038/s41592-019-0686-2

Vloeberghs, R., Urai, A. E., Desender, K., & Linderman, S. W. (2025). A Bayesian hierarchical model of trial-to-trial fluctuations in decision criterion. PLOS Computational Biology, 21(7), e1013291. 10.1371/journal.pcbi.1013291

Voss, A., Lerche, V., Mertens, U., & Voss, J. (2019). Sequential sampling models with variable boundaries and non-normal noise: A comparison of six models. Psychonomic Bulletin & Review, 26(3), 813–832. 10.3758/s13423-018-1560-4

White, H. (1982). Maximum Likelihood Estimation of Misspecified Models. Econometrica, 50(1), 1. 10.2307/1912526

Wiecki, T. V., Sofer, I., & Frank, M. J. (2013). HDDM: Hierarchical Bayesian estimation of the Drift-Diffusion Model in Python. Frontiers in Neuroinformatics, 7. 10.3389/fninf.2013.00014

